# Anti-Biofilm Potential of Nanonized Eugenol against *Pseudomonas aeruginosa*

**DOI:** 10.1101/2022.12.19.521144

**Authors:** Sourav Ghosh, Upasana Sett, Anabadya Pal, Sanchita Nandy, Susmita Nandy, Soumajit Chakrabarty, Abhijit Das, Pathikrit Bandopadhyay, Tarakdas Basu

## Abstract

This study dealt with nanonization of eugenol, a major phytochemical present in basil leaf, which has pharmacological potential as an anti-bacterial agent. Eugenol nanoparticle (ENP) was synthesized by simple ultrasonic cavitation method through emulsification of hydrophobic eugenol into hydrophilic gelatin. Thus, the nanonization process made the water-insoluble eugenol to water-soluble nano-eugenol, making the nano-form bioavailable. The average size of the ENPs was 20-30 nm. Entrapment efficiency of eugenol within gelatin cap was about 80% of the eugenol, that was used as precursor in the nanonization reaction. *In vitro* release of eugenol from gelatin cap was slow and sustained over a period of five days. The ENP had higher anti-biofilm potency than eugenol for both formation and eradication of biofilm, formed by clinically relevant pathogen *Pseudomonas aeruginosa*. Minimal biofilm inhibitory concentration and minimal biofilm eradication concentration of ENPs were 2.0 and 4.0 mM respectively. In addition, the measurement of *P. aeruginosa* biofilm biomass, biofilm pellicle formation, biofilm thickness, amount of biofilm-forming extra-polymeric substance, cell surface hydrophobicity, cell swarming and twitching efficiencies, cellular morphology and biofilm formation in catheter demonstrated that the anti-biofilm efficacy of nano-eugenol was 30-40% higher than that of bulk eugenol. Thus, ENP can be used as a potential drug against pneumonia, a chronic infection in lung caused by *P. aeruginosa*, which is difficult to treat with antibiotics, due to natural intrinsic resistance of biofilm-formed cells to most antibiotics. The overall actions of ENP have been presented in the figure 1.

**Highlights:**

- Nano-formulation of eugenol, an important phytochemical, by ultrasonic cavitation method, which was simple, time-saving, low-cost and eco-friendly.
- Nanonization made water-insoluble eugenol into water-soluble form with enhanced therapeutic efficacy.
- The eugenol nanoparticle (ENP) could inhibit formation of biofilm as well as facilitate eradication of pre-formed biofilm of *P. aeruginosa*.
- Biofilm formation was found to be prevented significantly on ENP-coated catheter.
- Nano-eugenol may be used as a potential drug against bacterial diseases, caused by pseudomonal biofilm, which are difficult to treat by antibiotics.
- Nano-formulated eugenol may also be used as an effective anti-fouling agent for biomedical devices like contact lens, pace-maker, materials for organ transplantation etc. to prevent bacterial colonization.

## 1. Introduction

Bacteria are known to alternate between two forms – free swimming unicellular planktonic and sessile multicellular community as biofilm (1). Bacterial biofilm is a well-organized cluster of cells that display an ability of adhesion on a surface and is surrounded by a self-produced extra polymeric substance (EPS) that acts as protective shield for the embedded cells. This shield facilitates persistent survival of the resident cells and protects them from various types of threats including predations, antimicrobials and any kind of diverse and harsh conditions (2). Thus, EPS of biofilm hinders the action of antibiotics to the resident bacteria and assist the bacteria to develop antibiotic resistance ability rapidly. According to World Health Organization (WHO), biofilm-mediated antibiotic resistance causes death of about 700,000 people per year worldwide (3). Amongst various bacterial strains, most encountered and clinically important biofilm-forming pathogens are *Acinetobacter baumannii, Staphylococcus aureus, Staphylococcus epidermis, Vibrio cholerae and Pseudomonas aeruginosa* (4).

**Figure 1:**
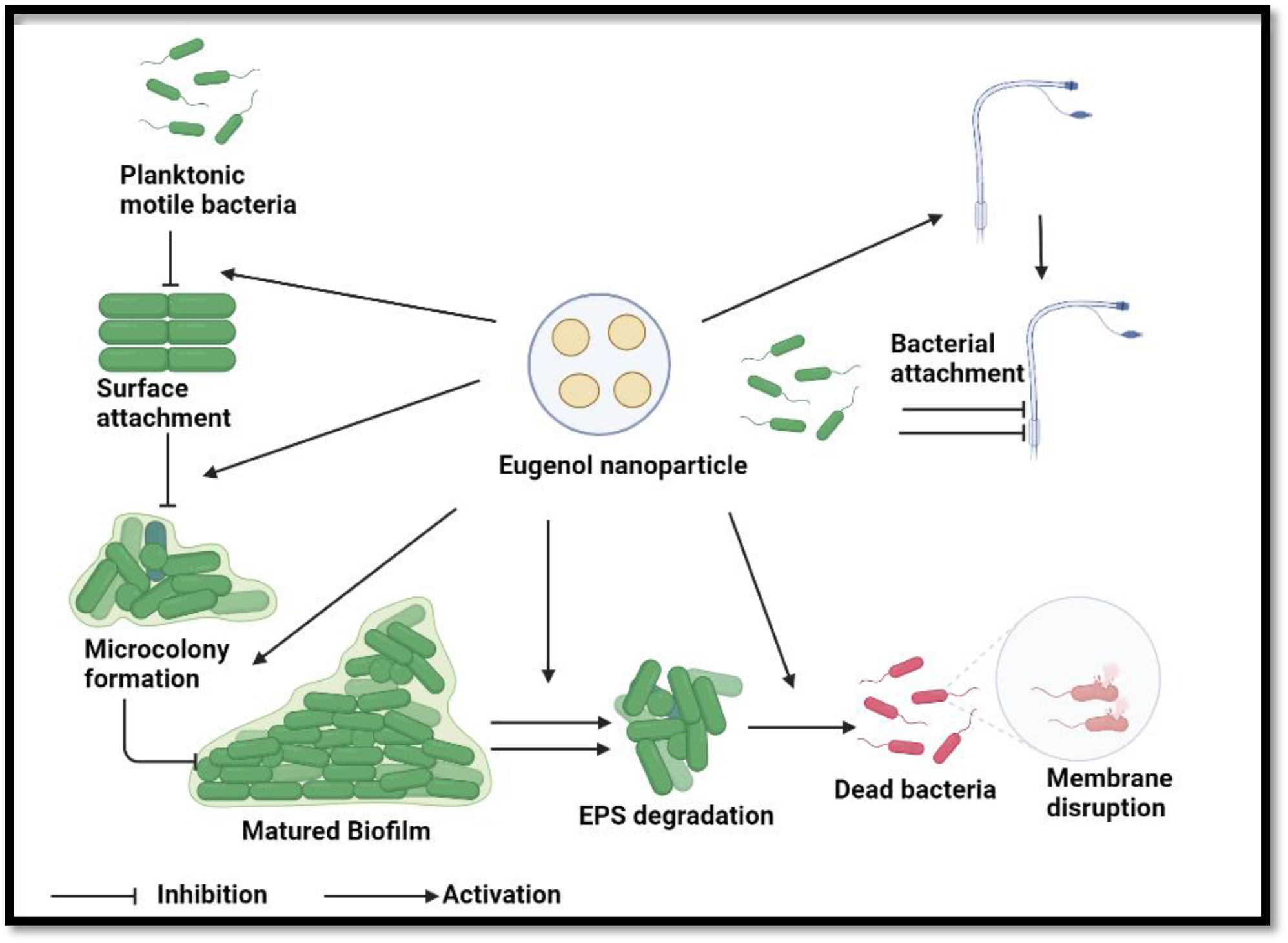
Graphical abstract showing overall anti-biofilm action of ENP that is capable to inhibit each and every step of biofilm formation. ENP also carries the potential to destroy EPS and kill the biofilm residing cells by membrane disruption. Bacterial colonization has been found to be prevented significantly on ENP-coated catheter.

The opportunistic pathogen *P. aeruginosa* causes complicated infections in the lungs of the individuals suffering from cystic fibrosis (CF), burn wound, cancer, external otitis, endocarditis, etc.; enabling chronic bacterial colonization on lung epithelial cells (5). This pathogen is also a cause of various nosocomial infections like bacteremia, urinary tract infection (UTI) and sepsis (6), generally transmitted from local infections such as pneumonia or burn wound into systemic blood stream, for which the global mortality rate is about 13-50% from country to country (7). This gram-negative pathogen creates a global fear of untreatable infections. The ‘WHO’ listed this pathogen as greatest public health threat (8). To address the issue of antibiotic inaction, researchers are exploring for development of effective anti-biofilm agents, which have one or multiple action(s) like inhibition of planktonic cell attachment to a site, killing of biofilm-forming planktonic cells, induction of preformed biofilm eradication factors (dispersin, alginate lyase), degradation of EPS, etc. In this regard, some natural antimicrobials like essential oils (thymol, carvacrol, eugenol), obtained from the plant’s secondary metabolites, have promising antibacterial /anti-biofilm activity against a wide range of gram-negative and gram-positive bacteria (9). Such agents of green origin have less side effects compared to any other conventional therapies (10). These metabolites have gained much interest as alternative of antibiotics for their broad-spectrum antibacterial potency, multiple mechanisms of action and low tendency to induce resistance (11). However, their potential clinical application has been greatly limited by their poor solubility in aqueous media and low stability. To overcome these problems, nanonization of such essential oils through entrapment into natural polymers have been designed, with the property of controlled release of the antimicrobials from the nanocarrier polymers. Such drug delivery nano-system has been demonstrated as promising antimicrobial weapons against bacterial biofilms [(12) -(13)].

In this study, we ventured to synthesize a nano-form of the phytochemical ‘eugenol’ - a phenylpropanoid, by entrapping within biocompatible gelatin molecule. The reasons behind selecting eugenol were – a) it has versatile pharmacological potentials as antimicrobial, antioxidant, anesthetic, analgesic, anticonvulsant, anticancer, neuro-protective, hepato-protective, cardio-protective, etc. (14) and b) it is a major component of cinnamon bark, clove bud, basil leaves etc., which are well available and therefore cheap in price in Indian market. The well-dispersed and stable eugenol-nanoparticles (ENPs), which we designed and prepared, was investigated to have promising anti-biofilm activity, compared to free eugenol, against clinically relevant pathogen *Pseudomonas aeruginosa*.

## 2. Materials and methods

### 2.1 Nanoformulation of eugenol

First of all, 9.8 ml aqueous solution of gelatin (3mg/ml) [Merck] was taken in a conical flask, that was placed on ice and was sonicated by a probe sonicator [Cole-Parmer, CPX 130] at 70% amplitude and 20 KHz for 10 min (in discontinuous mode; pulse on for 10 sec and pulse off for 10 sec) to obtain a homogenous solution of gelatin. To this gelatin solution, a volume of 200 µl eugenol (from a stock of 1g/ml) [Sigma-Aldrich] was added and the mixture was further sonicated on ice, as described above, for 10 min to obtain stable eugenol-gelatin nano-composite suspension of milky-white color. For purification of nanonized eugenol, the prepared suspension was centrifuged at 15000 rpm for 15 min, after which the pellet of eugenol nanoparticles was washed twice with and finally dispersed in 10 ml of Milli-Q water and stored at room temperature.

### 2.2 Characterization of Eugenol Nanoparticles (ENPs)

#### 2.2.1 Size, shape, PDI (poly-dispersive index) and zeta potential

The size and shape of the nanoparticles were measured by Atomic Force Microscopy (AFM), Field-Emission Scanning Electron Microscopy (FE-SEM), and Transmission Electron Microscopy (TEM). For AFM study, a smear of about 1000 times diluted ENPs was made by the process of drop casting on clean square cover slip (22 mm). The cover slip was scanned by AFM [Veeco, di-Innova] in tapping mode, using the nanoprobe cantilever made of silicon nitride and the obtained images were analyzed by Nanoscope software, as described in (15). For study with FESEM [F-50, FEI], separate cover slip was gold coated in ultra-vacuum condition on carbon tape (15). The images were analyzed by Image-J software. For TEM study, sample was prepared on carbon-coated copper grid [Applied Biosystems], as described in [16; (16) and the study was done in HR-TEM [JEOL, JEM-2010], operated at an accelerated voltage of 200 KV. The PDI and zeta potential of the synthesized ENPs were measured by dynamic light scattering (DLS) instrument [Malvern, Nano-ZS].

#### 2.2.2 Entrapment efficiency (EE) of eugenol within gelatin coat

Eugenol was reported to have an absorption peak around 280 nm (17). Therefore, the EE i.e., the percentage of entrapped eugenol into gelatin cap was estimated by measuring the (absorbance)_280nm_ of the total amount of eugenol that was used as precursor for the synthesis of ENPs and the (absorbance)_280nm_ of the residual amount of unentrapped eugenol in the centrifuged supernatant of the as prepared ENP suspension (as obtained during purification of the synthesized ENPs, described in subsection-2.1). The EE was finally calculated by following the equation –

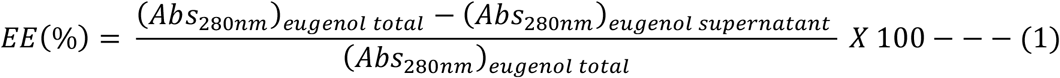

Since eugenol exhibits a concentration dependent antioxidant potential, the EE was also determined by measuring the antioxidant activity of the total amount of precursor eugenol and that of unentrapped eugenol in the centrifuged supernatant by the methods like DPPH assay and ABTS assay.

#### 2.2.2 DPPH radical scavenging assay (18)

An amount of 60 µM methanolic, purple color solution of free radical DPPH (1,1-diphenyl-2-picryl hydrazyl) [Sigma-Aldrich] was reduced by antioxidant like eugenol to yellow colored compound 1,1-diphenyl-2-picryl hydrazine. The decrease in absorbance of DPPH was recorded at 517 nm. Therefore, the antioxidant activity of drug (eugenol) was inversely proportional to OD_517 nm_ and accordingly, the percentage of entrapment i.e., entrapment efficiency (EE) of eugenol within gelatin was determined by the equation:

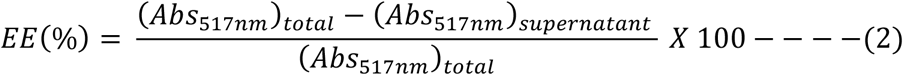

#### 2.2.2 B ABTS radical scavenging assay (19)

Here, a 7.0 mM aqueous solution of the radical cationic chromophore ABTS^.+^ (2,2’-azinobis-[3-ethylbenzothiazoline-6-sulphonic acid diammonium salt] [SRL] was mixed with equal volume of 2.45 mM potassium persulfate and was kept in dark for 12-14 hr, to obtain finally a stable blue-green colored solution of ABTS. The solution had absorption peak at 734 nm and the value of (absorbance)_734nm_ decreased with reduction of ABTS by any antioxidant and consequently the blue-green color faded. Here also, the formula for determination of EE was the same, as represented by the above equation-2, replacing the (absorbance)_517nm_ by (absorbance)_734nm_.

#### 2.2.3. Release of eugenol from ENPs

This parameter was measured by dialysis method (20). The purified ENP suspension (500 µl) was taken in a dialysis bag with 10 KDa cut-off membrane [Sigma-Aldrich] and the bag was immersed in a beaker containing 40 ml phosphate buffer saline (PBS - NaCl:137 mM, KCl: 2.7 mM, Na_2_HPO_4_:10 mM, KH_2_PO_4_:1.8 mM, pH-7.4). Dialysis was allowed to continue at 37^0^C. At different intervals of time (3, 6, 12, 24, 48, 72, 96 and 120) hr, an aliquot of 1.0 ml was withdrawn from the beaker and the volume was replenished by adding 1.0 ml fresh PBS. Concentration of released eugenol in the withdrawn aliquot was estimated by its absorbance at 280 nm.

### 2.3 Anti-biofilm activities of ENP

#### 2.3.1 Biofilm culture

To study the anti-biofilm activities of ENP, the rod-shaped, aerobic, pathogenic bacterium *Pseudomonas aeruginosa* MTCC 2488 (21) was selected as target organism. The bacterial cells were grown in nutrient broth [Himedia] at 37^0^C in a gyratory shaker [Labcompanion, IS-971R] at 125 rpm up to (OD)_600nm_ of 0.2 (∼10^8^ cells/ml). A volume of 0.1 ml grown culture was taken in each well of 96-well plate and allowed to grow further at 37^0^C for 24 hr without shaking, to allow biofilm formation. The nutrient broth medium was then removed and the wells were washed with PBS to remove non-adherent bacteria. The adhered cell film was analyzed to study the anti-biofilm role of nano-eugenol, from the following different aspects.

#### 2.3.2 Determination of minimal biofilm inhibitory concentration (MBIC) and minimal biofilm eradication concentration (MBEC) of an anti-biofilm agent

For a particular biofilm forming bacterium, MBIC (minimum biofilm inhibition concentration) of an anti-biofilm agent is defined as the lowest concentration which causes more than 90% of inhibition of biofilm formation, after 18-24 hr incubation of the planktonic bacterial cells with the agent, whereas MBEC (minimum biofilm eradication concentration) of an agent is defined as the lowest concentration which eradicates more than 90% of an established biofilm, when incubated with the agent for 18-24 hr (22). To determine the MBIC of an anti-biofilm agent, pseudomonal biofilms were allowed to form, as described in subsec. 2.3.1, in well plate in the presence of different concentrations of the agent. On the other hand, MBEC was determined by incubating pre-formed biofilm with different concentrations of antibiofilm agent at 37^0^C for 24 hr. The state of biofilm was analyzed by using blue-colored non-fluorescent dye resazurin; in presence of reducing environment of living cells, the dye turns into fluorescent pink color compound, which exhibits excitation and emission at 530 and 590 nm respectively [(23)-(24)]. In principle, the relative fluorescence unit (RFU) is proportional to the number of living cells and therefore, the extents of i) inhibition of biofilm formation and ii) biofilm eradication were estimated by measuring RFU. The biofilm was treated with 0.1ml resazurin sodium salt [Sigma-Aldrich], dissolved at a concentration of 0.1% (w/v) in PBS.

#### 2.3.3 Assessment of biofilm biomass

Biofilm mass was determined by staining the filmed cells with blue colored crystal violet (CV) dye. In principle, the intensity of blue color of film is directly proportional to biofilm biomass. In this study, biofilms were first washed with PBS to remove planktonic bacteria from well-plate and were then stained with 0.1% (w/v) aqueous solution of CV [SRL] for 30 min [(25)-(26)]. The stained biofilms were further washed with PBS to remove excess stain and were subsequently air dried for microscopic [Invitrogen, Evos FL auto] study. Thereafter, the films were allowed to solubilize by 95% ethanol and finally the absorbance of the solubilized sample was recorded by a microplate absorbance reader [Bio Rad, iMark™] at 595nm; biofilm biomass was quantitated from absorbance value.

#### 2.3.4 Assessment of biofilm pellicle formation

Pellicle is a kind of bacterial assemblage, formed majorly at air-liquid interface and minorly at solid-liquid interface. Pellicle is more clearly visible in glass tube than well plate culture (27). For assessing pellicle inhibition and eradication, same experimental method, as described in subsec. 2.3.3, was followed for staining cells with CV and subsequent imaging and spectroscopic studies for qualitative and quantitative measurements respectively.

#### 2.3.5 Assessment of biofilm thickness

Syto 9 is an excellent green fluorescent nucleic acid stain that is permeant to both prokaryotic and eukaryotic cells, having intact or damaged cell membranes and bind DNA, exhibiting fluorescence with excitation and emission at 483 and 503 nm respectively (28). In this study, the biofilm was allowed to grow on individual coverslips by putting them inside the wells of a 6-well plate and adding 6 ml of grown cells in the wells, for subsequent incubation at 37 ^0^C for 24 hr in static condition. The formed biofilm was stained with Syto 9 [Invitrogen]. Biofilm thickness measurement and 3D images were acquired from the fluorescence intensity through Z-stacking method (29) by ‘confocal laser scanning microscopy’ [Carl Zeiss, LSM 800].

#### 2.3.6 Assessment of biofilm forming extra-polymeric substance (EPS)

EPS, in biofilm architecture, is composed of cellular polysaccharide in major and DNA in minor amount. The dye Congo red (CR) binds EPS and the intensity of red color of the dye is directly proportional to the quantity of EPS (30). For qualitative measure of EPS, bacterial biofilm was stained with 0.1% (w/v) aqueous solution of CR [Himedia] for 45 min and subsequently microscopic study was performed. For quantitative estimation of the EPS, the CR-incorporated biofilm was solubilized in dimethyl sulfoxide (DMSO) [Merck] and the absorbance of the solubilized sample was recorded at 528 nm.

#### 2.3.7 Assessment of cell surface hydrophobicity (CSH)

CSH was assessed by using the principle of bacterial adhesion to non-polar hydrocarbon n-heptane [Spectrochem, India] (31). Bacterial planktonic cells were centrifuged and a cellular suspension in PBS was prepared which displayed optical density (OD) A_0_ in between 0.6 to 0.7 at 600 nm. The cell suspension was then overlaid by n-heptane in 8:1 (v/v) ratio, vortexed vigorously, phases were allowed to be separated for 10 min and the OD (A_1_) of aqueous phase was recorded at 600 nm. The percentage of hydrophobicity was measured by the equation:

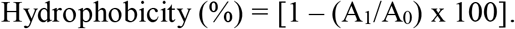

#### 2.3.8 Assessment of flagellar swarming motility

Swarming efficiency (SE) is totally dependent on flagellar functionality. Swarming occurs when cells interact with surface, reprogram their physiology to adapt biofilm features (32). SE was assessed on dry agar plates consisting of 0.8% nutrient broth, 0.5% agar [SRL] and 0.5% glucose [Merck]. A volume of 5 µl, having 10^8^ cells/ml, was inoculated on each plate and incubated overnight. SE was determined by measuring the diameter of zone of movement (by Vernier calipers) on agar plates.

#### 2.3.9 Assessment of bacterial type IV pili mediated twitching

Bacteria twitch on solid surfaces by successive extension and retraction of surface filaments known as type IV pili. This mechanotactic movement aids in surface attachment, the first step of biofilm formation (33). Twitching potential (TP) was assessed on agar plate containing 1% tryptone [Himedia], 0.5% yeast extract [Merck], 1% NaCl [Merck] and 1% agar [SRL]. The twitch plates were dried and a volume of 5 µl, having 10^8^ cells/ml, was inoculated at the center bottom of each plate by dipping the micro-tip through agar, and allowed to incubate for overnight. TP was measured from the diameter of zone of bacterial motility (by Vernier calipers).

#### 2.3.10 Assessment of bacterial morphology in biofilm

To observe the cellular morphology of pseudomonal biofilm, FESEM analysis was performed (34). Firstly, the bacterial cells were allowed to grow for 24 hr on coverslips, which were then washed and bacterial loads on them were fixed with 2% (w/v) aqueous solution of paraformaldehyde, followed by air-drying. The dried samples were subsequently gold coated on carbon tape under ultra-vacuum condition and finally imaged with FESEM.

#### 2.3.11 Assessment of bacterial colonization in catheter as biofilm

Implantation of catheter into the urinary bladder of patient’s body, suffering from prostate enlargement, epidural anesthetic, kidney damage, bladder fistula, etc.(35), is very common and *P. aeruginosa* readily causes nosocomial infections, forming biofilm on catheter’s surface inside. To study bacterial colonization in catheter, pieces (1.0 cm x 0.5 cm) of Foley’s urinary catheter segment were incubated with freshly grown culture of *P. aeruginosa* (∼10^8^ cells/ml) at 37^0^C for 24 hr. After incubation period, catheter pieces were aseptically recovered from bacterial culture and gently rinsed with PBS to remove unbound bacterial cells. The colonized bacterial cells in catheter pieces were then fixed with 2% (w/v) aqueous solution of paraformaldehyde [Merck] for 20 min. The catheter segments were then gold coated in ultra-vacuum condition on carbon tape and imaged with FESEM.

#### 2.3.12 Statistical analysis

All the experiments were done thrice. Quantitative data are presented as mean ± standard deviation. Statistical significance of the results was determined by one-way analysis of variance (ANOVA) with Tukey’s multiple comparisons test, using Graph Pad Prism 9 software. P values, represented by different number of stars (*), demonstrate the significance level of experimental results as: ns-not significant; * - <0.01; ** - <0.001; ***-<0.0001 and **** - <0.00001.

## 3. Results

### 3.1 Synthesis and characterization of ENP

Eugenol nanoparticles (ENPs) were prepared by simple ultrasonic cavitation method (Fig:2), using eugenol and gelatin as the two cheap and well-available precursors. The method of preparation took less than 1.0 hr to be completed. Our following experimental results unravel that ENP represents an effective anti-biofilm agent, which is easy, time-saving to prepare and economically beneficial also.

**Figure 2:**
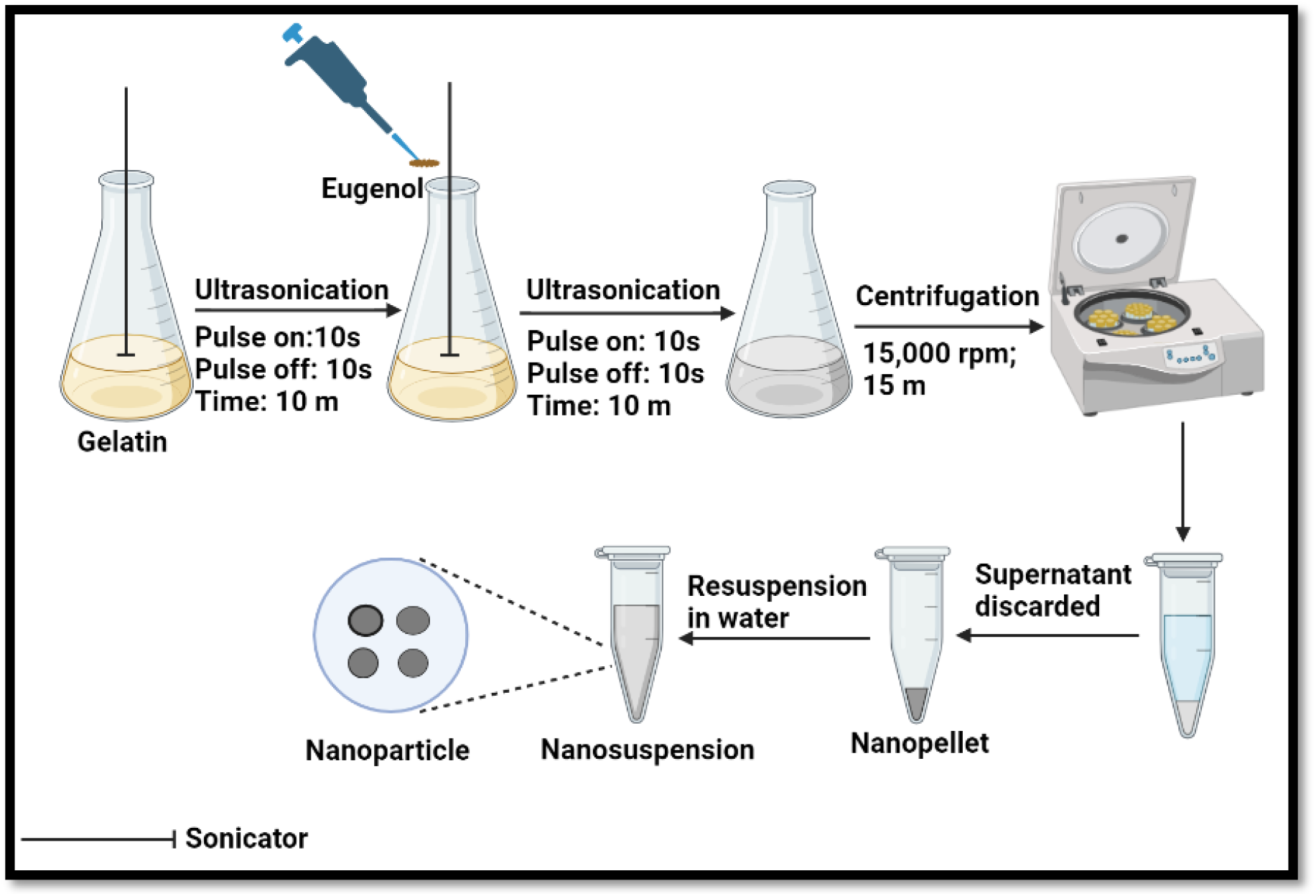
Method of preparation of ENP. ENP was synthesised by ultrasonic cavitation method and each single particle contains 4-5 dark spots.

The size of the prepared ENPs was measured to be (28 ± 12) nm by AFM (Fig. 3A), (20 ± 5) nm by FESEM (Fig. 3B) and (20 ± 8) nm by TEM (Fig 3C) i.e., the average size of the particles may be claimed as 20-30 nm. Such particle size is small enough than the diameters of numerous drug administration channels (efflux pump, ion channels, transporter protein) in bacterial cells (36) and so, ENPs have high chances of translocation into the bacterial cells. Moreover, AFM, FESEM and TEM images depicted that the particles were spherical in shape. Furthermore, TEM image (Fig 3C, marked by circles) exhibited that, each particle consists of 3-4 dark spots that can be assumed as eugenol molecules are adsorbed on the surface of gelatin globule. The value of poly-dispersive-index (PDI) of the ENP suspension, as measured by DLS-instrument was (0.29 ± 0.2). Such PDI value signified that the particles were very much monodispersed, because according to International Standard Organizations (ISOs), PDI value ≤ 0.5 depicted mono-dispersed (equal sized) particles whereas PDI value ≥ 0.7 meant poly-dispersed (different sized) particles (37). Mono-dispersed nature of ENPs was also evident from single, sharp peak of the (particles’ size distribution vs. intensity) plot (Fig. 3D) obtained from DLS result. In addition to the PDI value, fig.3D shows that the DLS-measured size of the ENPs were (372 ± 7) nm, which was much larger than the average size of the same particles (20-30 nm), as determined by AFM, FESEM and TEM; the reason might be that DLS generally measures particles’ hydrodynamic size and in most cases this size appears to be more than that of the core particles and this is due to the presence of a hydration layer around a NP (38). The value of the Zeta potential [(17.92 ± 0.47) mV] of ENPs (Fig. 3E), as measured by the same DLS instrument implied that the particles were moderately stable. In principle, zeta potential (Z) of a suspending particle is defined as the electric potential developed on the particle’s surface between its surface charge and a thin layer of attracting counter ions (called stern layer) present in the suspending medium. Therefore, ‘Z’ causes the electrostatic repulsion between the adjacent particles in a dispersion medium (39); higher magnitude of ‘Z’ (negative or positive) corresponds to higher magnitude of repulsive forces between the particles and thereby higher stability of the particles preventing aggregation/flocculation. Particles having zeta potentials > ± 30 mV are normally considered stable (40) and so the ENPs can be claimed to have moderate stability.

**Figure 3:**
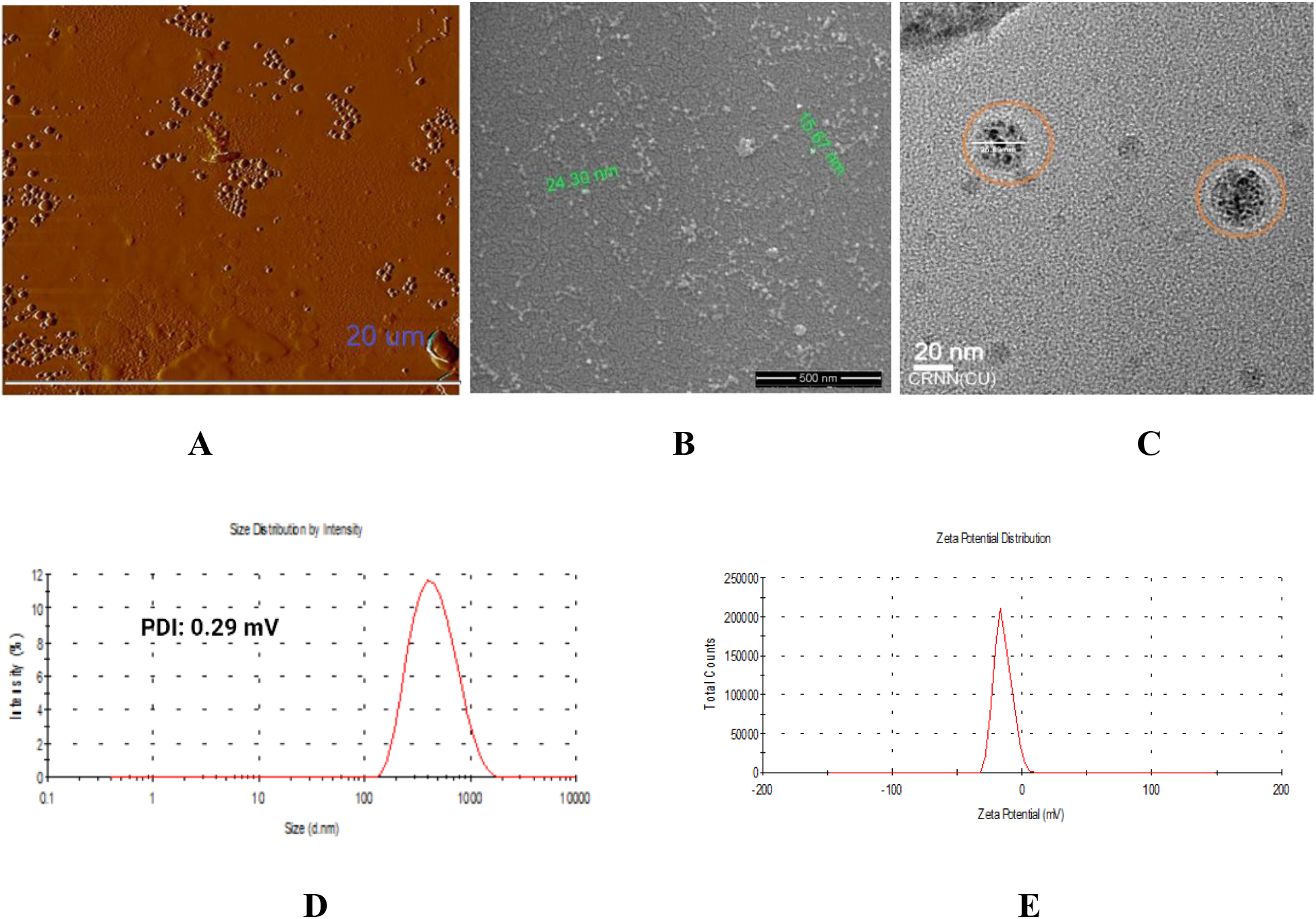
Size, shape, PDI and zeta potential of ENPs. A: AFM image; B: FESEM image; C: TEM image of nanoparticles; D: DLS-measured PDI from particles’ size distribution spectrum and E: DLS-measured zeta potential spectrum.

Entrapment efficiency (EE) of eugenol within gelatin cap was determined to be respectively (84.06 ± 2.1), (82.5 ± 1.25) and (77.37 ± 2.5)% from direct measurement of (Absorbance)_280nm_ of eugenol as well as indirect measurement of antioxidant activity of (Fig.4), as obtained from the three different experiments, were close to each other and depicted that about 80% of the total eugenol (20 mg/ml), used as a precursor in the nanonization reaction (mentioned in subsec. 2.1), was nanonized by our preparation method and thus the amount of eugenol per ml of the final centrifuged stock of ENP in 10 ml Milli-Q water was 16 mg/ml. In vitro release of eugenol from gelatin cap in PBS medium, as examined by dialysis technique, was found to take place over a period of five days and the release profile depicted slow and sustained release of the compound at physiological pH 7.4. About a half of the entrapped eugenol was released fast in first 24h and the residual 50% of eugenol was released slowly in the next 96h (Fig. 5). Thus, the high antibacterial efficacy of our ENPs, as discussed below, might be considered to originate from the initial bulk release of eugenol that killed invading bacteria within 24 hr of administration, followed by subsequent slow release of eugenol that maintained the killing environment for much longer period of about 4-days at target site.

**Figure 4:**
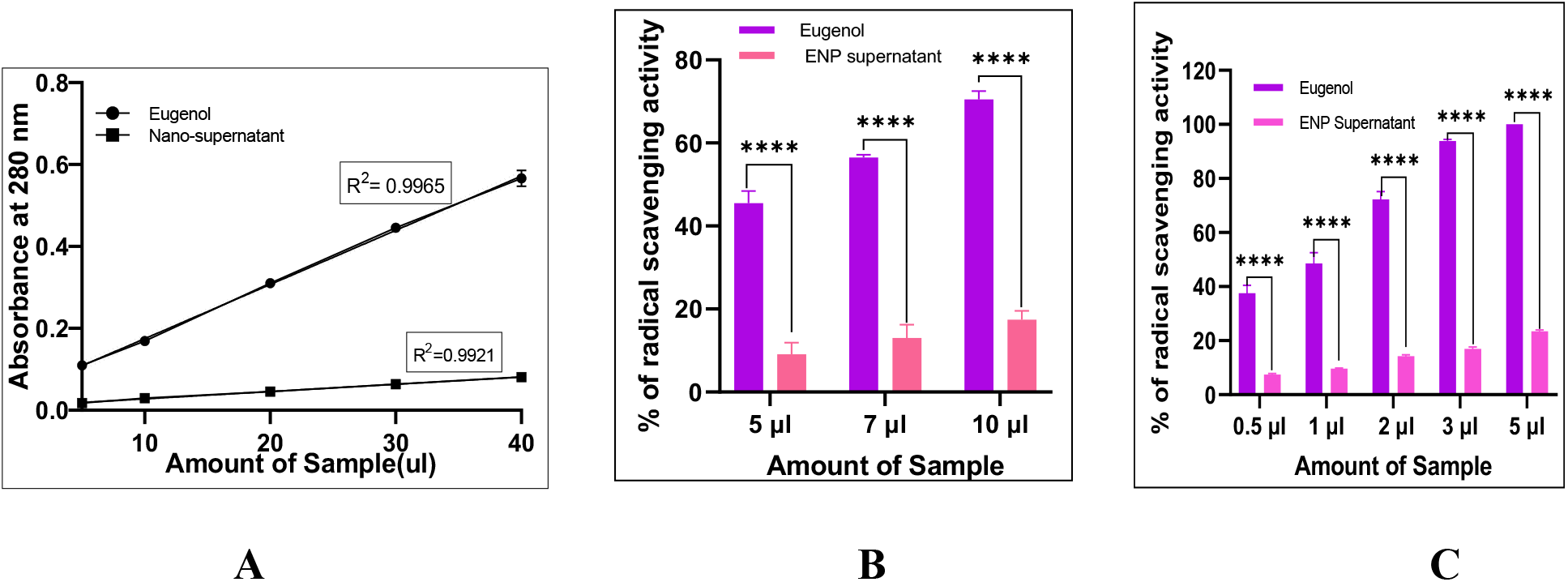
Entrapment efficiency of eugenol within gelatin cap. A: spectrophotometric assay of eugenol absorbance; B: DPPH and C: ABTS assays of eugenol antioxidant activity. For 4A, a solution of eugenol was prepared at concentration of 1µg/µl and the absorbance were taken at the corresponding points while nanosupertanat (obtained during preparation of nanoformulation) was diluted 20 times and the absorbance were taken.

**Figure 5:**
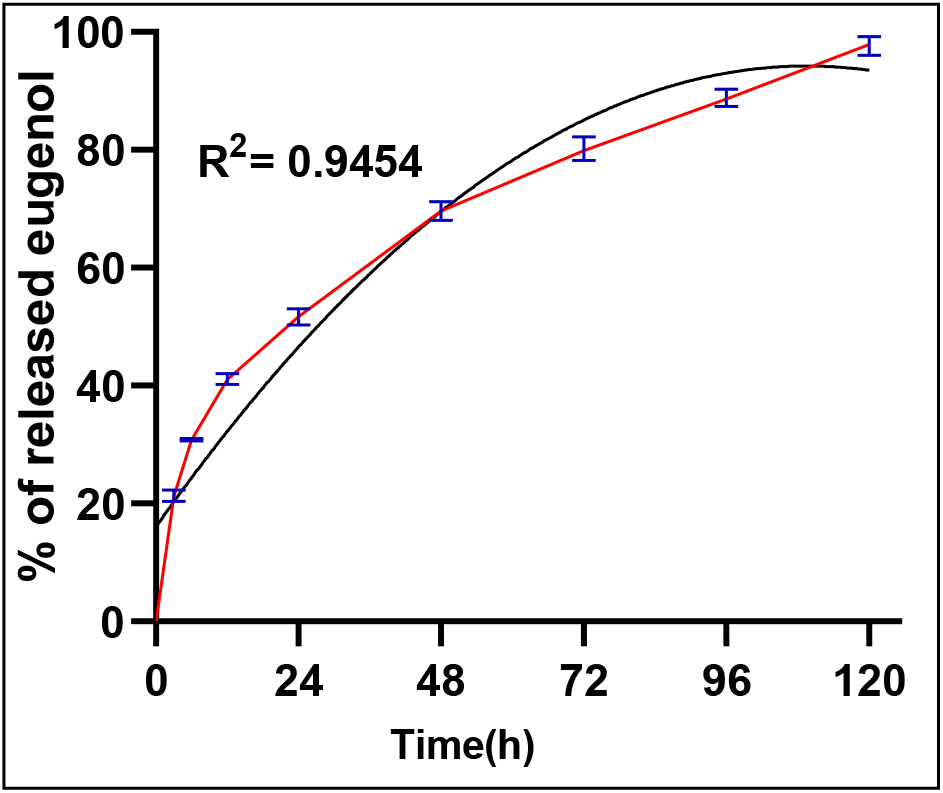
Release profile of eugenol from ENP. Blue bar represents the percentage of released eugenol at the corresponding time point whereas black line is non-linear fit of curve i.e. second order polynomial fit.

### 3.2 Anti-biofilm role of ENP

MBIC and MBEC doses of ENP were first determined as described in subsec. 2.3.2. For MBIC determination, when the concentration of ENP was increased from (0 - 2.5) mM, more than 90% inhibition of biofilm formation (in terms of bacterial viability) occurred at about 2.0 mM concentration, signifying that the MBIC value of ENP was 2.0 mM; similarly, when the concentration of ENP was increased from (0 - 4.5) mM, more than 90% eradication of preformed biofilm occurred at 4.0 mM concentration, signifying that the MBEC value of ENP was about 4.0 mM (Fig. 6). It should be mentioned here that the molar concentration of ENP meant the molar concentration of the eugenol, that was entrapped within the gelatin coat (i.e. 16.0 mg/ml, as discussed in the previous subsec. 3.1). With respect to the MBIC and MBEC doses of ENP, equivalent concentration of eugenol (2.0 and 4.0 mM respectively) was found to have the percentage of inhibition and eradication as about 67.5 and 69.5 respectively (Fig. 6). However, eugenol itself was water insoluble and its nanonization through entrapment within water-soluble gelatin made it soluble in water and so, bioavailable. Therefore, to study the effect of bulk eugenol, it was dissolved in methanol and so, while the anti-biofilm efficacy of bulk eugenol was checked, control experiment was also performed with equivalent concentration of methanol itself, that was carried over with eugenol. Similarly, when the antibiofilm efficacy of ENP was studied, control experiment was also performed with equivalent concentration of gelatin only that was carried over with eugenol. The results depicted that methanol and gelatin individually had inhibited biofilm formation by about 17.0 and 2.0% respectively and had eradicated pre-formed biofilm by about 21.0 and 1.0% respectively (Fig. 6). Since the anti-biofilm effects (inhibition and eradication both) of methanol with respect to eugenol was considerably high, after subtraction of the contribution of methanol, the real biofilm inhibition and eradication percentage of 2.0 and 4.0 mM of free eugenol was about 50 and 48% respectively. The anti-biofilm effect of gelatin only, with respect to ENP, by 1-2% could be considered negligible. It can, therefore, be summarized as – the anti-biofilm effect of nano-eugenol was about 45-47% more than that of bulk eugenol.

**Figure 6:**
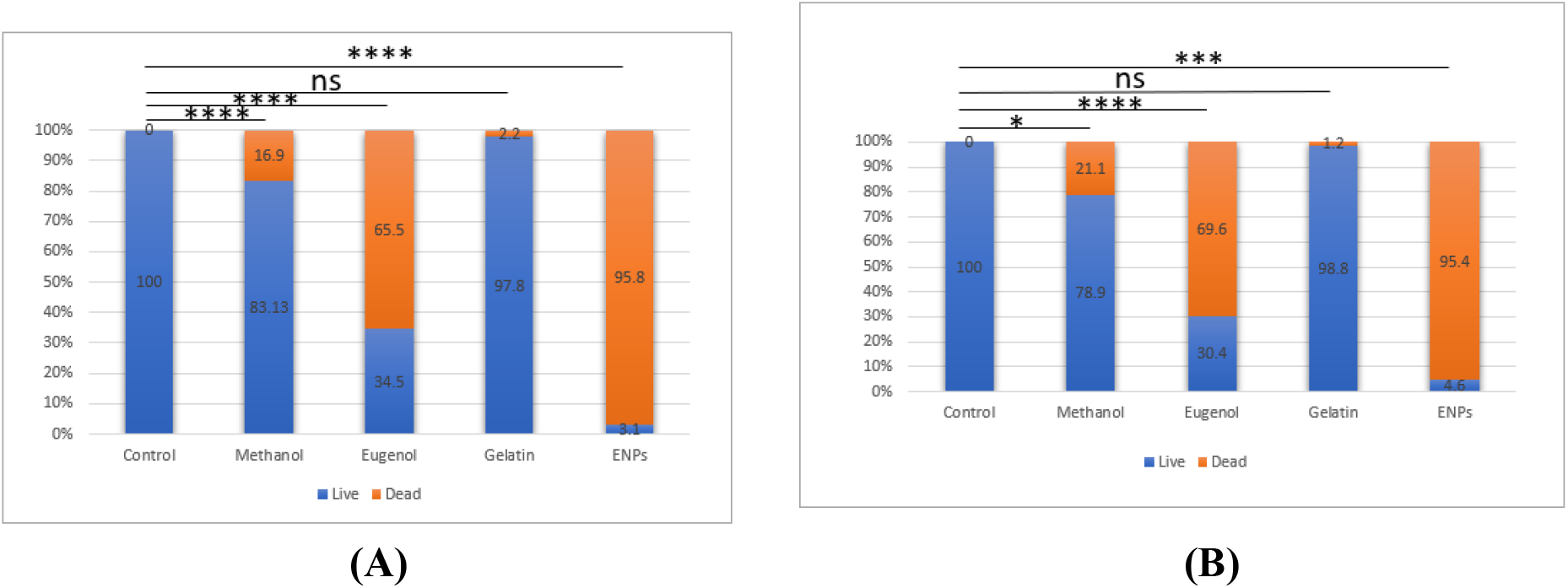
Percentage of biofilm inhibition (A) and eradication (B) in terms of dead cell.

Biofilm biomass is defined as the total mass of surface attached bacteria and their protective covering EPS (41). At MBIC and MBEC doses of ENP, estimation of biofilm biomass was performed by the method of CV-staining, as described in subsec. 2.3.3. The qualitative result at MBIC, as represented by the microscopic images of CV-stained biofilm, has been depicted in Figs. 7(A-E); the quantitative result, as represented from the spectroscopic measurement of (Absorbance)_595nm_ of dissolved stained-biofilm, is depicted in Fig. 7F; the corresponding qualitative and quantitative results at MBEC have been shown in Figs. 7(G-K) and Fig. 7L respectively. It is clearly observed from the Figs. 7(A-E) & 7(G-K) that the relative biomass of formed and eradicated biofilms (due to treatment with MBIC [2.0 mM] and MBEC [4.0 mM] doses of ENP and equivalent concentration of eugenol, methanol, gelatin, and no agent [untreated control] separately), as measured by the CV-staining intensity (I) of biofilms was in the descending order as (BM)_Control_ > (BM)_Gelatin_ > (BM)_methanol_ > (BM)_Eugenol_ > (BM)_ENPs_. The quantitative result, represented by fig. 5F shows that during biofilm formation, the biomass of formed films in presence of ENP, eugenol, methanol, and gelatin was respectively (82 ± 0.5), (67 ± 3.7), (16.5 ± 3.9) and (1.46 ± 1)% less than the biomass of biofilm formed by untreated control cells; whereas, Fig. 5L illustrates that during biofilm eradication, separate presence of above agents reduced film biomass by (92 ± 3.6)_ENP_, (67 ± 2.5)_Eugenol_, (17.9 ± 2.7)_Methanol_ and (3.6 ± 1.6)_Gelatin_% from mass of the film formed by control cells. Thus with respect to biofilm biomass, a) the inhibitory capacity towards biofilm formed and b) the eradication capacity of pre-formed biofilm by ENP were respectively about 30 and 40% more effective than that of free eugenol (real capacities of eugenol was calculated by subtracting the contribution of methanol).

**Figure 7:**
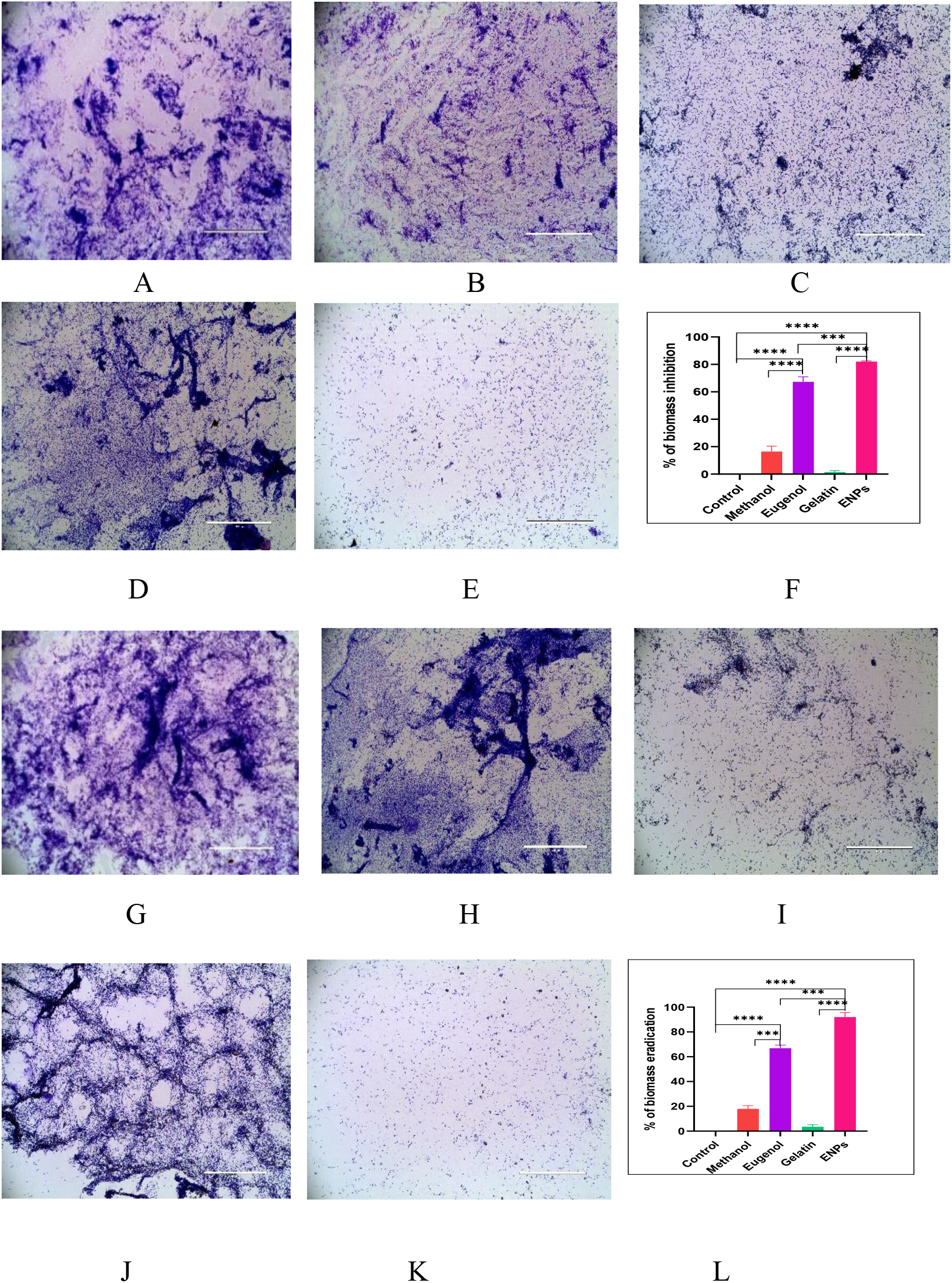
Decrease of biofilm biomass with biofilm inhibition (A-F) and eradication (G-L). CV-stained biofilms, A: Control; B: Methanol; C: Eugenol; D: Gelatin; E: ENP; F: Quantitative representation of biomass during biofilm formation and G: Control; H: Methanol; I: Eugenol; J: Gelatin; K: ENPs; L: Quantitative representation of biomass during biofilm eradication. White bar on each figure represents 200 µm

Pellicle is another form of microbial assemblage (biofilm), formed at interface regions of culture tubes – mostly at air-liquid interface as a prominent intense pellicle ring and also at glass-liquid interface. The qualitative results, as obtained from microscopic study (as described in subsec.2.3.4) on inhibition of pellicle formation and eradication of pre-formed pellicle by ENP, eugenol, methanol, gelatin, and no agent (untreated control) separately have been presented in Fig.8(A-E) and Fig.6(G-K) respectively; the quantitative results, as obtained from spetrophotometric study, have been presented in Figs.6F and 6L. The MBIC and MBEC doses of ENP caused inhibition of pellicle formation as well as eradication of preformed pellicle by about 90.0 % while equivalent doses of bulk eugenol, methanol and gelatin exhibited inhibition by (72.4 ± 3.15), 34.4 and 19% respectively, and eradication by (69.3 ± 0.8), 23.8 and 7.36% respectively. Thus with respect to pellicle formation or eradication, the effect of MBIC and MBEC doses of ENP was found to be about 50 and 40% higher than that of the equivalent concentrations of bulk eugenol (after subtracting the contribution of methanol effect, the real effect of eugenol itself was found out).

**Figure 8:**
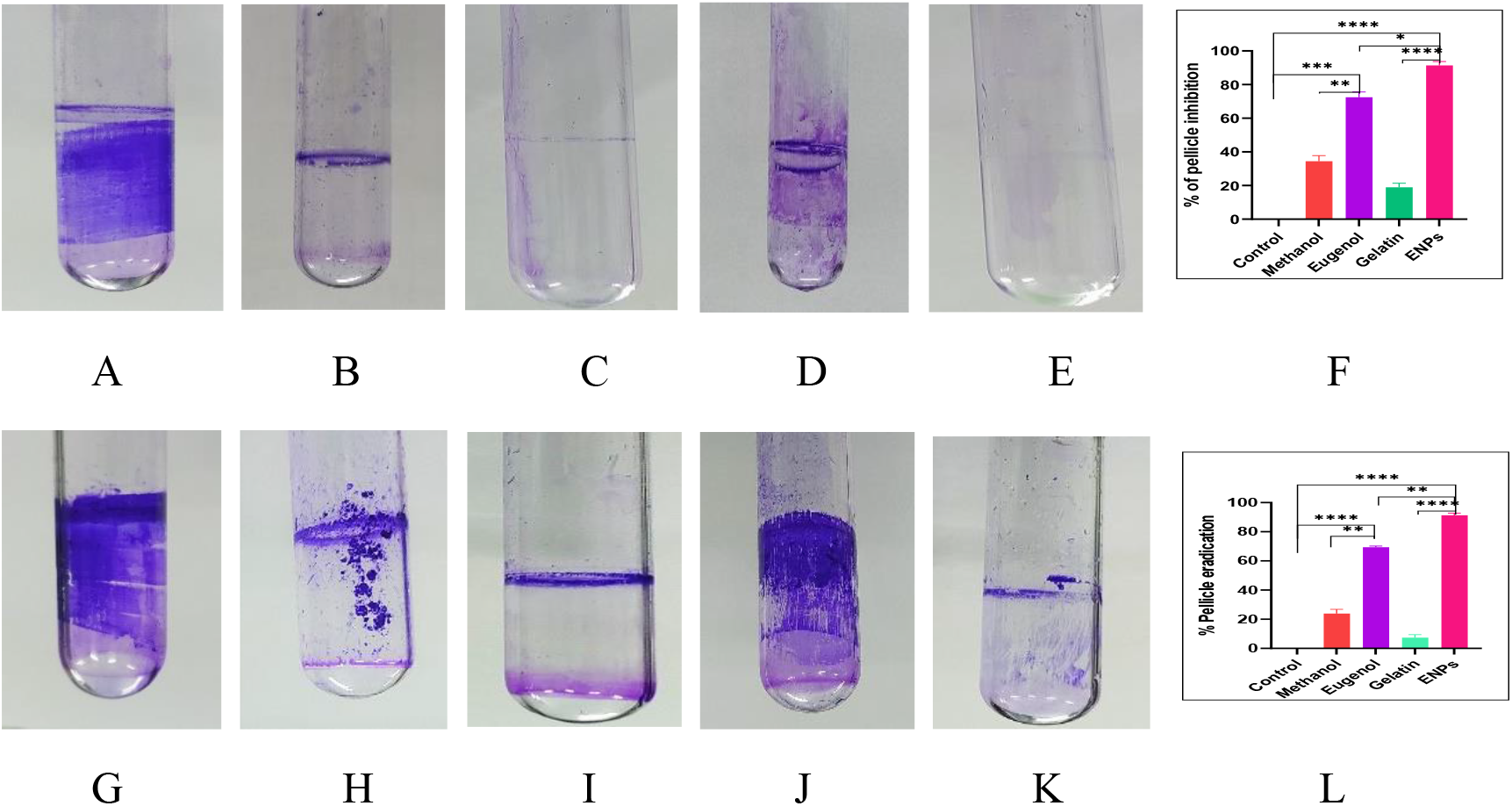
Decrease of pellicle formation (A-F) and increase of pellicle eradication (G-L). CV-stained pellicles, A: Control; B: Methanol; C: Eugenol; D: Gelatin; E: ENP; F: Quantitative representation of pellicle inhibition during its formation and G: Control; H: Methanol; I: Eugenol; J: Gelatin; K: ENP; L: Quantitative representation of pellicle removal during biofilm eradication.

Biofilm growth and thickening occurs in both horizontally and vertically by addition of new cells on pre-existing cells in overlapping manner (42). The qualitative and quantitative results on biofilm thickness a) during inhibition of biofilm formation by cell growth in presence of 2.0 mM ENP (MBIC) and 2.0 mM free eugenol separately, at 37^0^C for 24 hr and b) during eradication of pre-formed biofilm by its incubation with 4.0 mM ENP (MBIC) and 4.0 mM free eugenol separately, at 37^0^C for 24 hr, have been presented in Fig. 7. The extent of fluorescence intensity of the confocal microscopic images (Fig. 9[A-C] and Fig. 9[E-G]) implied qualitatively that during biofilm formation and eradication both, in presence of nano- and bulk eugenol, film thickness followed descending order as (thickness)_control cells_ > (thickness)_bulk eugenol treated cells_ > (thickness)_nano eugenol treated cells_. Figs. 9D and H exhibit quantitatively that thickness of biofilm decreased down by a) about 59 and 43 % respectively, during biofilm formation, in presence of 2.0 mM of each of ENP and free eugenol separately, and b) about 73 and 48 % respectively, during biofilm eradication, in presence of 4.0 mM of each of ENP and free eugenol separately, compared to the thickness of the biofilm formed by control cells. Thus, this result was in consonance with those of biofilm biomass and biofilm pellicle formation and eradication.

**Figure 9:**
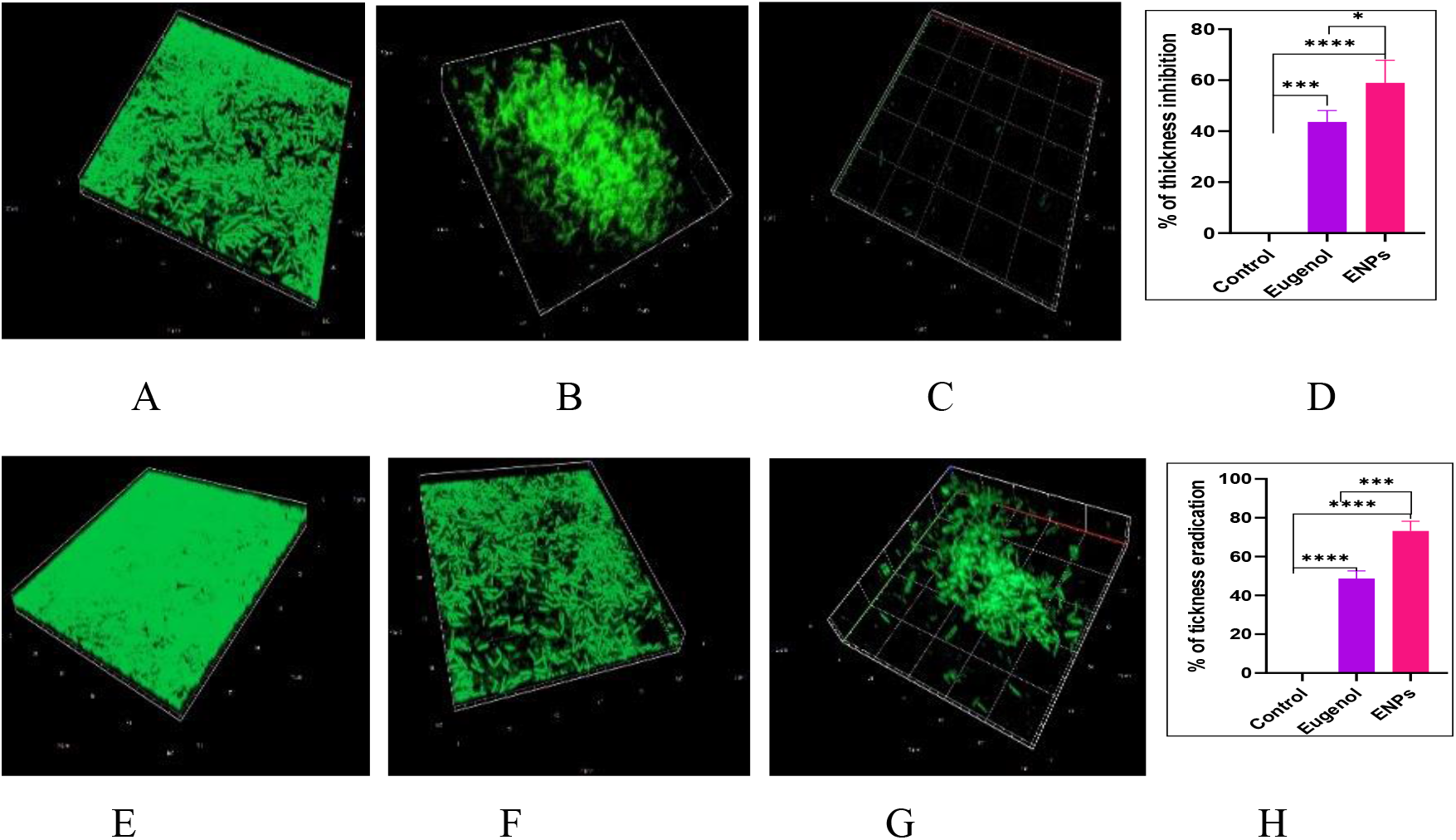
Lowering of biofilm thickness with inhibition (A-D) and eradication (E-H). Syto 9-stained biofilms, A: Control; B: Eugenol; C: ENP; D: Percentage of lowering of film-thickness, during biofilm formation and E: Control; F: Eugenol; G: ENP; H: Percentage of lowering of film-thickness, during biofilm eradication.

Pseudomonal EPS contains majorly three exopolysaccharides like alginate, Psl and Pel, together with extracellular DNA (eDNA) in minor amount (43). CR staining of biofilm, followed by microscopy qualitatively demonstrated more reduction of EPS amount, during both biofilm formation and eradication, by the administration of respectively 2.0 mM and 4.0 mM ENP than the same molarities of eugenol (Fig. 10[A-E] and [G-K]). The spectrophotometric study on the solubilized CR-incorporated biofilm in DMSO, as described in subsec. 2.3.6 quantitatively depicted that a) during biofilm formation, ENP and free eugenol separately caused reduction of EPS amount by about 92 and 65 % respectively, whereas b) during biofilm eradication, ENP and free eugenol caused reduction of EPS amount by about 99 and 82%, compared to the amounts of control biofilm (Fig. 10D & G). The reduction in case of methanol and gelatin were 20 and 7.0 % respectively (during biofilm formation) and 31 and 10 % respectively (during biofilm eradication). Therefore, with respect to EPS reduction, ENP was about 47-48% more efficient than bulk eugenol in both biofilm formation and eradication. The transformation of bacteria from planktonic to biofilm state is primarily initiated by surface attachment. Such attachment is dependent upon complex interactions among bacteria, solid substratum and the liquid phase (culture medium). The interactions are influenced by the involvement of the factors like cell surface hydrophobicity (CSH), flagellar motility (swarming), cellular twitching (extension and retraction of surface appendage type IV pili) etc. [(44)-(45)-(46)]. Therefore, the role of ENP and equivalent amount of eugenol, methanol, gelatin and no agent (untreated control) separately, on the above three factors of biofilm formation was also investigated. Experimental results illustrate that a) CSH was reduced by about 93, 79, 64 and 49% (Fig. 11a), b) bacterial swarming was retarded by about 83, 39, 25 and 5 % (Fig. 11b) and c) pseudomonal twitching got reduced by about 99, 51, 34 and 10%, by nano-eugenol, bulk-eugenol, methanol and gelatin respectively (Fig. 11c), compared to values in case of untreated control cells. Thus, ENP was more effective than free eugenol to reduce CSH by about 78%, to inhibit swarming by about 69% and bacterial twitching by about 82%.

**Figure 10:**
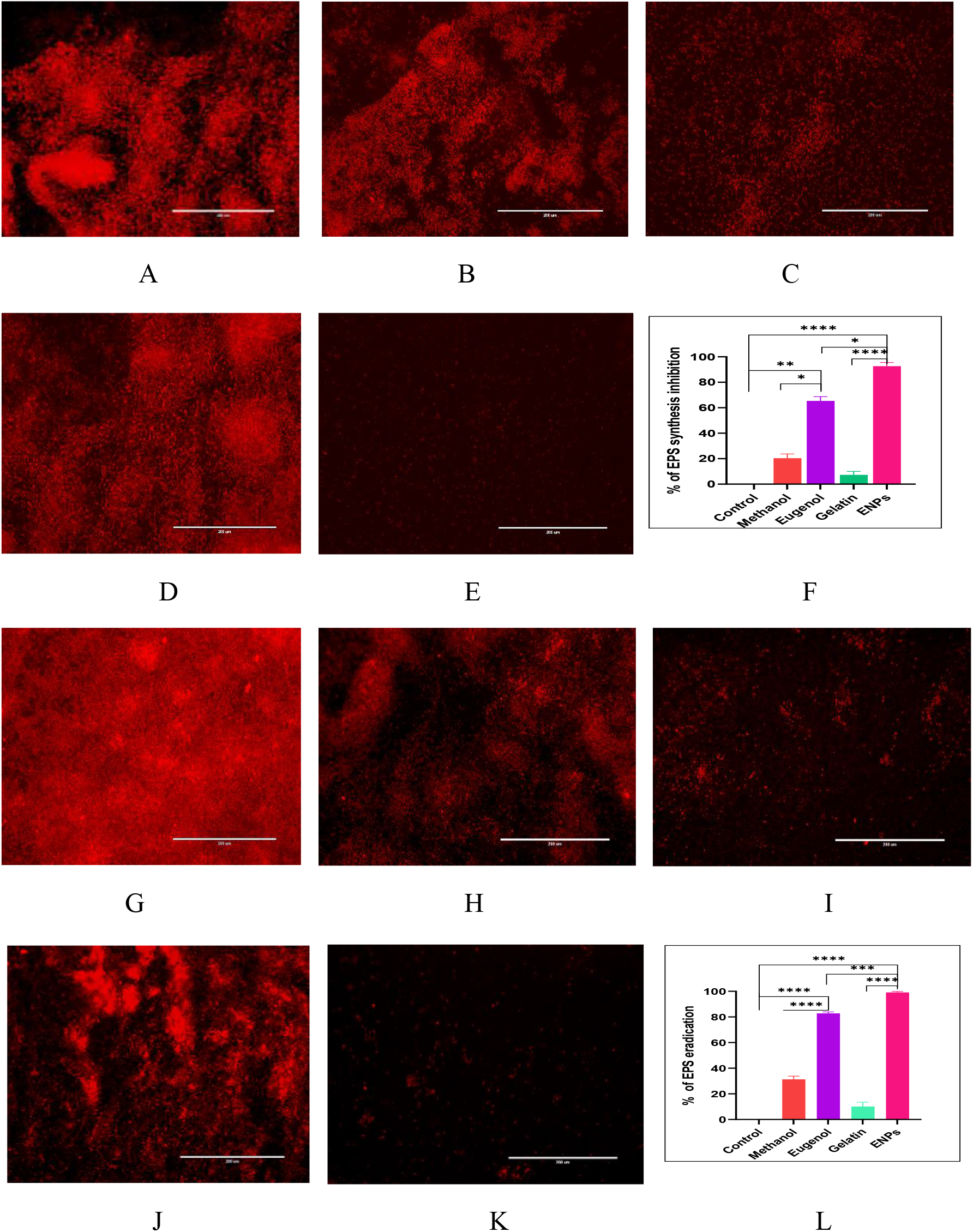
Reduction of EPS amount with inhibition of biofilm formation (A-F) and biofilm eradication (G-L). CR-stained biofilms, A: Control; B: Methanol; C: Eugenol; D: Gelatin; E: ENPs; F: Percentage of EPS reduction, during biofilm formation and G: Control; H: Methanol; I: Eugenol; J: Gelatin; K: ENPs; L: Percentage of EPS reduction, during biofilm eradication. White bar on each figure represents 200 µm

**Figure 11:**
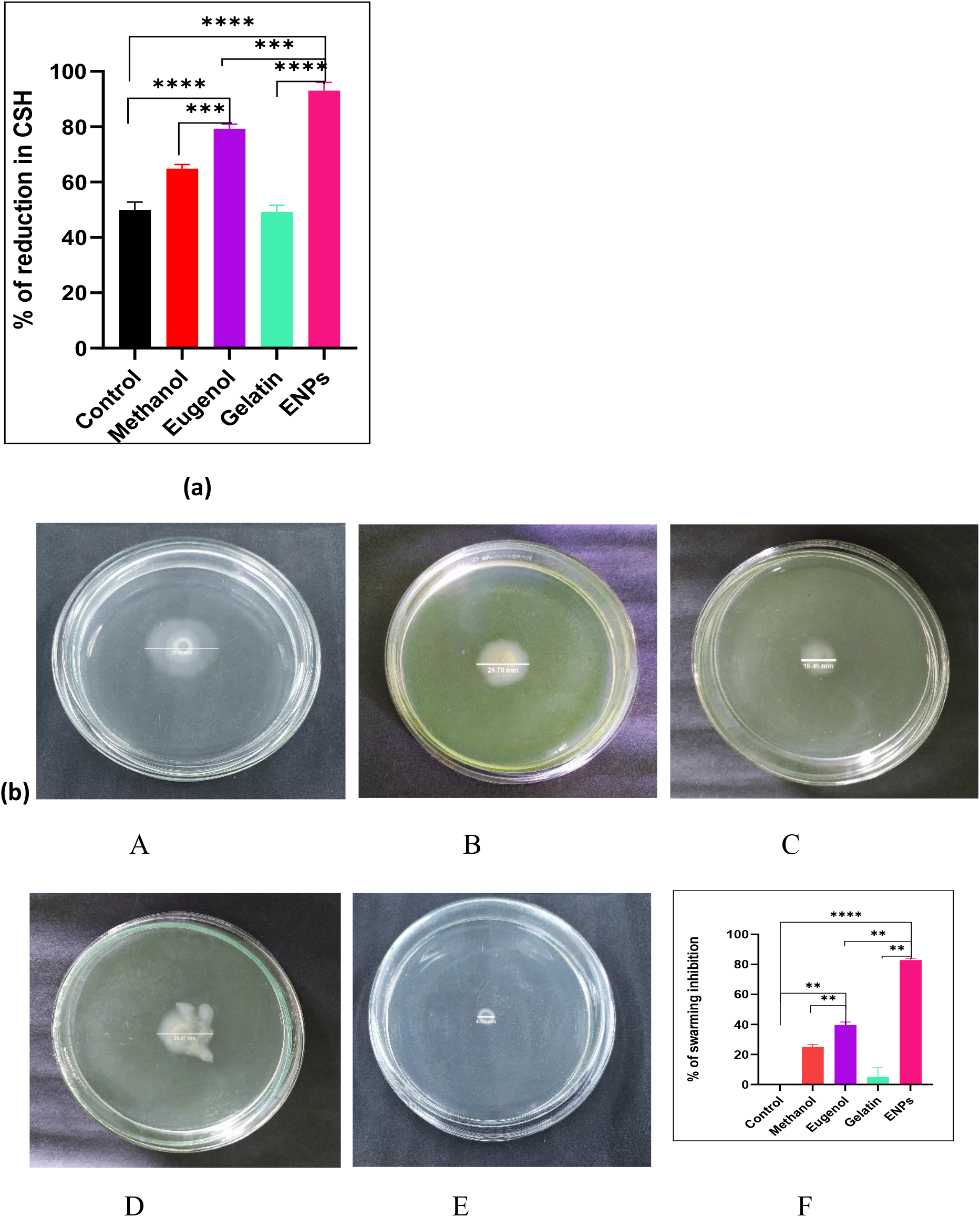
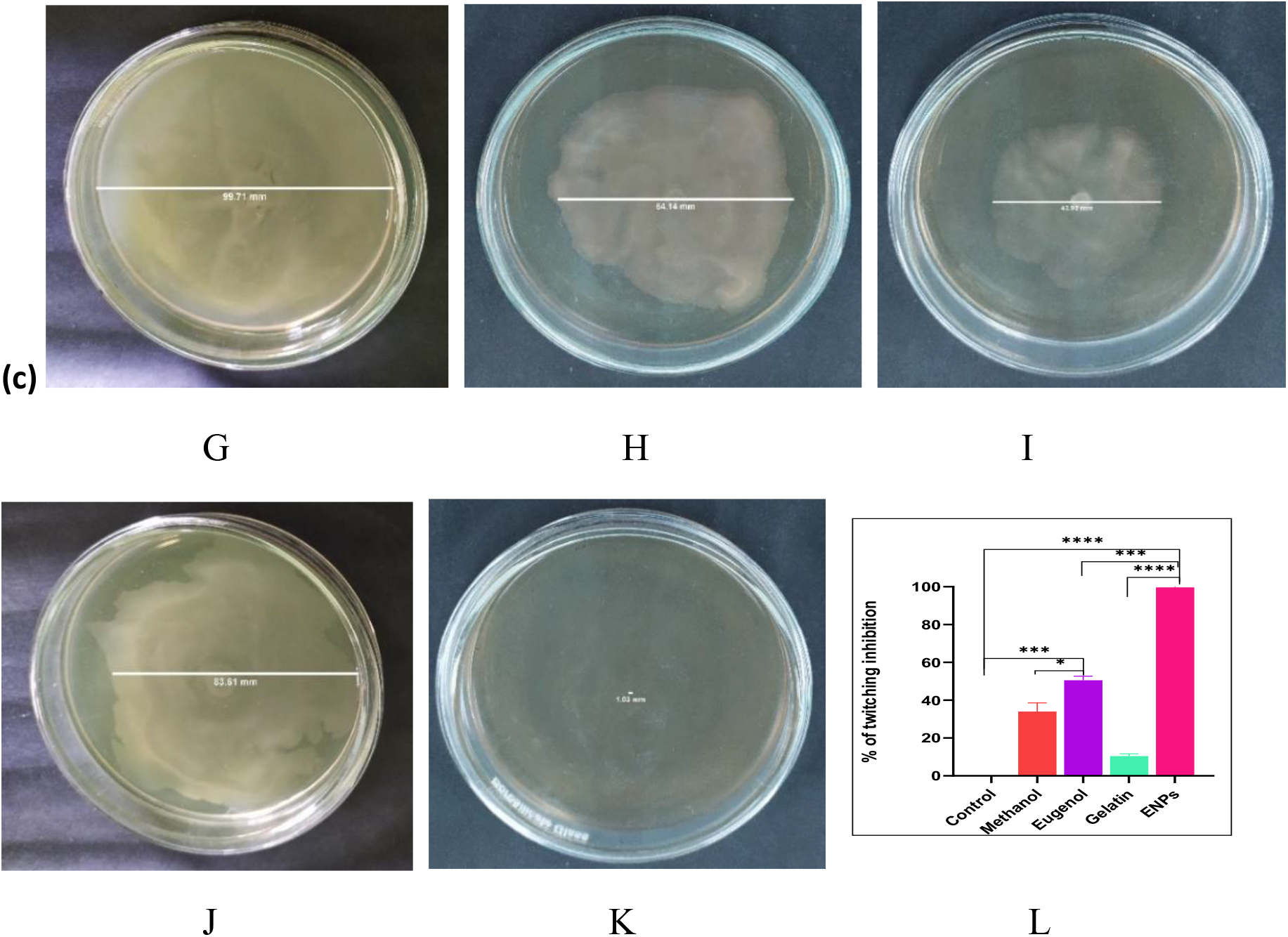
Percentage of (a) CSH decrease, (b) Swarming inhibition, and (c) Twitching inhibition, during biofilm formation. A: Control; B: Methanol; C: Eugenol; D: Gelatin; E: ENP and F: Quantitative representation of swarming during biofilm formation; G: Control; H: Methanol; I: Eugenol; J: Gelatin; K: ENP and L: Quantitative representation of inhibition of twitching during biofilm formation.

When cellular morphologies were studied through FESEM at MBIC dose of ENP (2.0 mM) and free eugenol (2.0 mM) separately, i) under no treatment condition, cells exhibited rod-shaped morphology with smooth outer lining (Fig.12A) that attributes perfect structural cellular integrity, ii) the eugenol-treated cells, some appeared as blistered rod-shaped structure along with some protrusions on cell membrane and some as spheroplasts (cells without cell wall) (Fig.12B) and iii) cellular morphologies of the ENP-exposed cells were completely disrupted with appearance of cellular fragments (fig12C).

**Figure 12:**
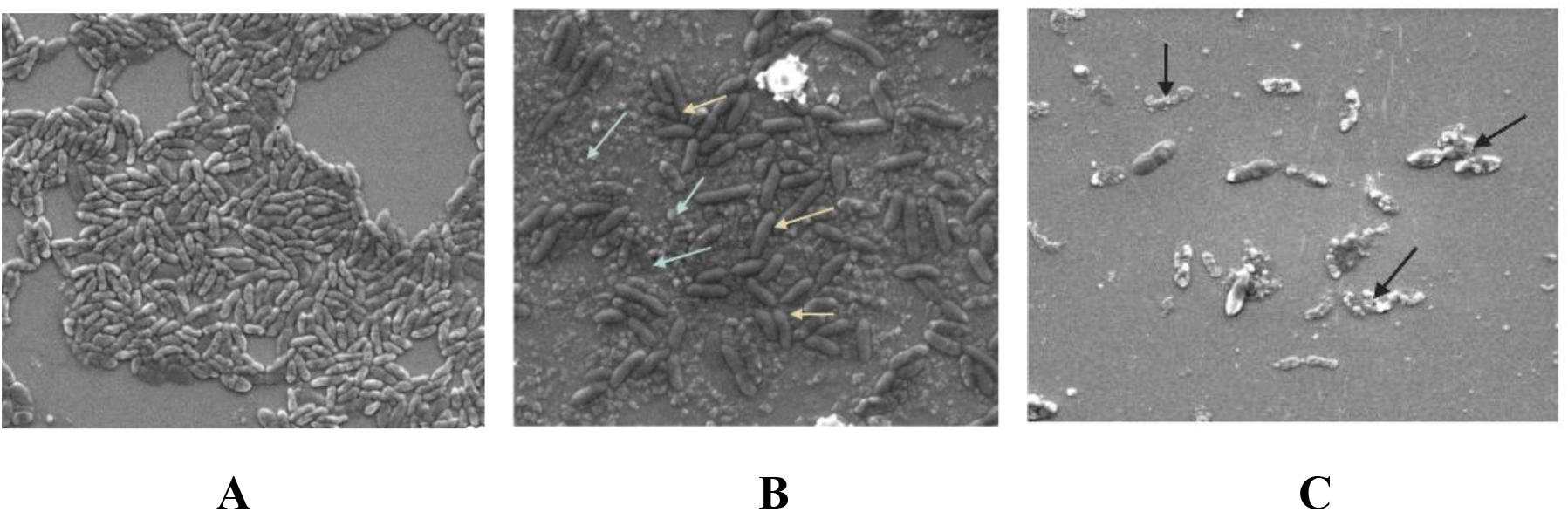
Cellular morphologies during biofilm formation. A: Control; B: Eugenol; C: green-white arrow indicates spheroplast, yellow arrow directs membrane blebbing and black arrow depicts cellular fragments

Pseudomonal biofilms are also formed on the surface of medical devices such as surgical instrument, dental material, contact lens, hip implant, pacemaker, catheter etc. Such bacterial aggregate is responsible for 60-70% healthcare-associated infections and is the major source of catheter-associated urinary tract infections (47). Several antibiotic-coated catheters are available, however these are literally ineffective against such biofilms of bacterial pathogens, as the films are hard to penetrate by antibiotics (48). Some other treatment strategies to prevent bacterial colonization on catheters had been reviewed using antiseptic, nitric oxide, silver, ammonium compound, chitosan, antimicrobial peptides and enzymes coating (47). We also ventured to develop green catheter by coating its inner surface with ENP. When the formation of biofilm was investigated on urinary Foley’s catheter under FESEM, i) Figs. 13A and 13B depicted respectively the images of bacteria-free smooth surface and thick biofilm-formed surface (by allowing bacterial growth, as described in subsec. 2.3.10) of uncoated catheter segments, ii) when catheter segment was coated with eugenol only, few planktonic cells were attached on its surface (Fig 13C), and iii) on ENP-coated surface, negligible number of cells were found to be attached (Fig. 13D). This study demonstrated that the ENP has great potential to be used as a green coating agent over surgical and biomedical instruments against biofilm formation by pathogenic bacteria.

**Figure 13:**
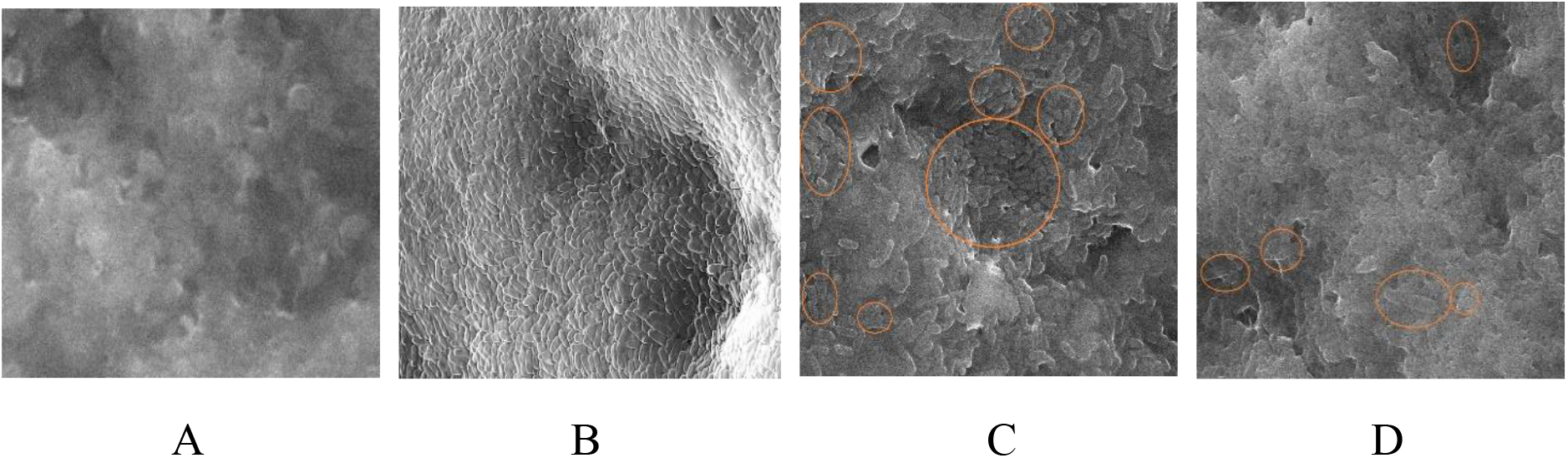
Role of ENP on bacterial colonisation in urinary Foley’s catheter surface. A: Bacteria-free smooth surface, B: Thick biofilm-formed surface, C: Eugenol-coated surface and D: ENP-coated surface of catheter segments. Circles in Fig C and D represent bacterial cells

## 4. Discussion

A potent nano-compound ENP was successfully synthesized by easy and simple ultrasonic cavitation method through emulsification of hydrophobic eugenol into hydrophilic gelatin. Synthesized ENPs were small enough in size, highly monodispersed, moderately stable and also have the capacity to hold a considerable amount of eugenol. ENP had a concentration dependent antibacterial potency on *P. aeruginosa in vitro* and at 2.0 mM concentration inhibited more than 90% of the bacterial biofilm formation, whereas at 4.0 mM concentration eradicated more than 90% of pre-established biofilm. Moreover, ENPs exhibited an initial bulk release and subsequent slow release of eugenol. Therefore, ENP could be contemplated as a potential drug candidate. Furthermore, ENP was found to be non-toxic in preliminary results (data not shown) on HEK 293T cell line *in vitro* and Balb/c mice *in vivo* at experimental doses.

Biofilm formation is a well-known phenomenon and occurs in multiple steps like attachment, micro-colony formation, biofilm growth and finally dispersion to progress the infection in another site. The transformation of bacteria from planktonic (freely suspended) to biofilm (surface attached) state is dependent upon complex interactions among bacteria, solid substratum and the liquid phase (culture medium). Those interactions are influenced by the involvement of biological factors (presence of particular surface protein, extracellular polymer, flagellar motility and CSH) of bacteria and the physicochemical factors (surface free energy, hydrophobicity, electrostatic charge) of the substratum, and the liquid phase (44). Flagellin, the major subunit of flagella and the flagellar cap protein function as adhesin to promote surface attachment (45). Eugenol significantly retards bacterial attachment and co-adhesion by down regulation of flagellar genes (*flaA, flaB and flgA*) encoding proteins and cell surface modifying gene (*waaF*) in *Campylobacter jejuni* (46). The second step of biofilm establishment i.e., the micro-colony formation is characterized by bacterial adhesion, aggregation and persistence; each of these phenomena are dependent on cell surface hydrophobicity (CSH). Generally, *P. aeruginosa* exhibits hydrophobic cell surface, which is correlated with slow growth, strong biofilm formation and persistence (49); clinical isolates of mucoid, slime-producing strains of *P. aeruginosa* become non-adherent by alteration of CSH, while non-mucoid, non-slime producing strain become adherent with high affinity by increasing CSH (50). Eugenol is shown to inhibit biofilm formation and adhesion on Hep-2 (human larynx epidermoid carcinoma) cell line and on polystyrene by decreasing of CSH in *Candida dubiliniensis and C. tropicalis* (51). The third step is biofilm growth, when bacterial cells are added on the pre-existing cells in an overlapping manner and such addition of new cells is facilitated by the presence of extra-polymeric substance (EPS). Eugenol is reported to reduce EPS production in *Vibrio parahaemolyticus* (12). Therefore, it is suggested that eugenol inhibited biofilm formation in each and every step of its establishment and our experimental results show that eugenol in nano-form to be more effective than bulk eugenol to inhibit biofilm generation capability in *P. aeruginosa*.

In pre-established biofilm, residing cells are protected by EPS barrier so that biofilm eradication is quite challenging for any antimicrobial agent. When pseudomonal cells attach and grow in sufficient number, cell-cell communication (quorum sensing) is activated to coordinate EPS production and cellular aggregation (52). During dispersion, some of the residing cells come out by breaking EPS barrier and form micro-colonies in other sites to spread the infection. In case of ENP-mediated eradication, biofilm liberated cells were found to be dead by the action of ENP and so, the infection progression was stopped. Pseudomonal EPS contains majorly three exopolysaccharides (namely alginate, Psl and Pel) and eDNA (extracellular DNA) in minor amount (43). Anionic alginate plays an important role in maintenance of biofilm architecture by providing adhesiveness and also helps in water retention in the film (53). During biofilm dispersion, *P. aeruginosa* produce alginate lyase enzyme which cleaves polysaccharide into short units (oligosaccharide) to promote detachment of bacteria away from surfaces (53). Eugenol and other essential oils are reported to have some characteristic interactive points for alginate, as analyzed by FTIR study [(54)-(55)] and perhaps through such interaction eugenol and ENP cause greater detachment of the bacteriofilm. The other two EPS exopolysaccharides, Pel and Psl function as primary matrix structural polysaccharide. The cationic polysaccharide Pel contains partially acetylated 1-4 glycosidic linkage of N-acetylgalactosamine and N-acetylglucosamine; it serves as primary scaffold for biofilm structure, elevates cell-cell aggregation leading to pellicle formation, cell attachment on the surfaces; whereas the neutral charged Psl, composed of repeating pentamers of D-mannose, D-glucose and L-rhamnose, is exclusively crucial for cell-surface attachment (56). The release of eDNA takes place by multiple pathways like cell lysis, quorum sensing, and pyocyanin-mediated membrane damage (57). Pel is also known to crosslink eDNA to provide more strength to EPS (56). Our results illustrated that the ENP caused reduction of biofilm thickness through interruption of cell-cell aggregation, which possibly occurred due to interaction of negatively charged ENPs with cationic Pel, rather than neutral charged Psl. Similar reduction of biofilm thickness with less activity of bulk eugenol than ENP signifies that interaction could also occur between alkaline Pel (due to presence of amine group) and weakly acidic eugenol. Significant EPS eradication and weak DNA binding affinity of eugenol (58) made it apparent that the configuration and complexity of major polysaccharides (Pel and or Psl) of *P. aeruginosa* were deformed by the action of eugenol. From our experimental results, it is suggested that i) both bulk- and nano-eugenol disoriented EPS structure and attacked the biofilm residing cells to kill, so that they could not be converted to planktonic state to progress infections, as seen in biofilm dispersion, and ii) a major number of cells were detached away from the surface as dead by the cleavage of alginate. Further studies are still required in more molecular detail to unravel the eugenol- and ENP-mediated biofilm eradication, especially the mechanism of EPS eradication in *P. aeruginosa*.

There are reports on the antibacterial mode of action of eugenol over a range of bacteria including *Salmonella typhi* (59), *Escherichia coli* (60), *Proteus mirabilis* (61), *Staphylococcus aureus* (62), etc.; but in all cases, the conclusive feature was only membrane disruption. In the published reports till date, it is concluded that eugenol exhibits multiple non-specific mode of action to retard bacterial infections. Firstly, eugenol breaks down the unsaturated fatty acid of bacterial membrane and thereby alter the membrane fatty acid profile of *Pseudomonas fluorescence, Salmonella enterica, Brochotrix thermosphacta*, etc.(63). Such kind of altered membrane facilitated loss of membrane integrity, as characterized by the leakage of intracellular content (potassium ion and ATP) into the exterior. Moreover, eugenol is also reported to downregulate the activity of the enzymes like histidine decarboxylase, amylase, ATPase etc.(64). Downregulation of ATPase is reflected by ATP leakage into the cell exterior, inhibition of ion transport and membrane mediated energy production. Membrane-coupled energy production is too much necessary for cellular structural integrity (that is governed by membrane integrity majorly) and cellular recovery under any impairment (65). It is suggested from our SEM analysis that eugenol could permeabilize and penetrate the outer membrane of *P. aeruginosa* by altering fatty acid profile; moreover, insertion of eugenol into inner membrane made it leaky and formed blistered membranous structure with some spheroplasts. ENP perhaps provided more interaction sites of eugenol and their action on both outer and inner membranes caused disruption of cellular membrane and thereby cellular structure. The antibacterial action of eugenol is, therefore, attributed as: alteration of fatty acid profile in outer membrane → destabilization of inner membrane → membrane blistering → bubble protrusion → spheroplast → burst cells.

## Acknowledgement

We are indebted to (1) UGC, for supporting the first author (SG) with his own research fellowship and contingency grant, (2) CSIR, for supporting the second and seventh authors (US and AD) with own respective research fellowship and contingency grant, (3) University of Kalyani, for supporting the third author (AP) with university research scholarship and contingency grant, (4) RUSA, for supporting the fourth author (SN) with project assistantship and partial support through the recurring grant under research project no. RUSA (C-10) / IP / 218, (5) DBT-GOI, for supporting the fifth author (SN) with research fellowship and partial support through the recurring grant under research project no. BT/PR28288/NNT/28/1558/2018U, (6) DST-GOI, for supporting the sixth author (SC) with his own INSPIRE research fellowship and contingency grant, (7) UGC-DAE for partial support through the recurring grant of its project no. CRS/2021-22/02/534, (8) DST-GOI for its “FIST” [SR/FST/LSI-623/2014(C)] and “PURSE” [SR/PURSE Phase 2/37(G)] Programs and UGC-GOI for its DRS(II)-SAP [F.5-3/2018/DRS-II(SAPII)], for providing different instrumental and infrastructural supports, and (9) Prof. Sukhen Das, Dept. of Physics, Jadavpur University, Kolkata for providing us the FESEM facility.

## Patent

This work has been patented (filed and published) through Indian IPR (Intellectual Property Right) under rules of regulations of government of India. Patent no. 202231052844 A; International Classification: A61L0029080000, C07K0014210000, A61K0031705600, A61L0029160000, A61K0009140000.

